# Rethinking Alzheimer’s: Novel miRNAs Illuminate a Disease Beyond the Brain

**DOI:** 10.1101/2025.11.27.690929

**Authors:** JS Novotný, M Čarná, EB Dammer, Z Mao, GB Stokin

**Author notes:** These authors contributed equally to this work (JS Novotný, M Čarná). **Corresponding author:** Gorazd B. Stokin, Institute of Molecular and Translational Medicine, Faculty of Medicine and Dentistry, Palacký University Olomouc, Hněvotínská 1333/5, 779 00 Olomouc, Czech Republic.

## Abstract

Alzheimer’s disease (AD) poses major health, social and economic challenges to the modern world. Despite the advances in understanding AD, our knowledge about its pathogenesis remains incomplete. Recent data suggest that circulating microRNAs (miRNAs) undergo complex changes in AD. Since these changes are yet to be comprehensively characterized, we investigated miRNAs in the context of AD using two meta-analytical approaches. We reproducibly identified 2895 miRNAs in a cohort of 4186 individuals from 22 studies. Here we show that 194 miRNAs exhibited widespread changes in AD, including some novel miRNAs not yet linked to AD. These novel AD miRNAs broaden the landscape of research on the role of miRNAs in AD. Targets of these miRNAs further uncovered many biological pathways that, to date, remain poorly understood in AD with several “AD miRNAs” never described in the brain. “AD miRNAs” described outside the brain significantly influenced interleukin signaling, Toll receptor signaling, p38 MAPK pathway and insulin/IGF pathway. Our results reveal a greater complexity of biological pathways involved in AD than previously thought and raise the question of whether AD is indeed a brain-specific and not a systemic disorder. These findings advance current understanding of AD pathogenesis and lay the ground for the development of next-generation AD biomarkers and design of miRNA-engaged therapies.

## Introduction

Short non-coding RNAs with antisense complementarity, known as microRNAs (miRNAs), were originally discovered to post-transcriptionally regulate gene expression during roundworm development^1,2^. Some miRNAs contain sequence codes (EXOmotifs) that are instructive of their secretion^3,4^ either packaged into vesicles or bound to proteins^5,6^. Secreted miRNAs were identified in all major body fluids^7^. By circulating, they allow for long-distance gene regulation and coordinated crosstalk between tissues^8^.

Abnormalities in miRNAs have been described in many human disorders from cancer^9^ to Alzheimer’s disease (AD)^10^. AD is the most common neurodegenerative disorder characterized by abnormal levels or processing of the amyloid precursor protein (APP) leading to amyloid plaques and aberrant phosphorylation of the microtubule-associated protein tau^11^. Dysregulated microtubule-dependent transport, mediated by molecular motors, underlies axonal transport impairments and pathology in AD^12^. Several other molecules and processes have been linked to AD pathogenesis including ApoE ε4 genotype^13^, the insulin pathway^13^ and past infections^14^. Accumulating evidence also suggests a degree of mutual exclusivity between development of cancer and AD^15,16^. Changes in miRNAs have been reported in the brain^17^, cerebrospinal fluid^18^ and plasma^19,20^ in AD. Recent work showed that abnormalities in miRNAs take place already during physiological aging and that miRNA changes in AD represent at least in part a pathological exacerbation of physiological aging^21^. We here examined systematically circulating miRNAs in AD using two meta-analytical approaches and rigorous data selection. We found that miRNAs changed in AD, including a number of novel miRNAs not yet linked to AD, target many biological pathways that are largely unaccounted for in the context of AD with several of them never described in the brain. These findings reveal greater complexity of biological processes involved in AD than previously thought and raise the question of whether AD is not a systemic rather than a brain disorder.

## Methods

### Study design

The study followed PRISMA 2020 reporting guidelines^22^. Study resources for meta-analysis were obtained by searching PubMed, PubMed Central (PMC) and Gene Expression Omnibus (GEO) repository using keywords [*miRNA AND (AD OR alzheimer) AND (plasma OR serum OR blood)*]. All studies reported to investigate human circulating miRNAs in AD until October 1, 2024, that provided either complete raw miRNA expression data or miRNA differential expression (DE) data with fold-changes and P-values for all analyzed miRNAs (significant and non-significant miRNA changes) were included in the meta-analysis. Research involving non-human subjects, studies not reporting original research, research of miRNAs isolated from sources other than plasma, serum or blood, reports without full text, experiments presented by incomplete results, for example experiments reporting only miRNAs with significant raw or differential expression data, but no information about miRNAs with non-significant changes, and contributions presenting insufficient and/or missing data were excluded.

### Data extraction

Three researchers conducted literature search, retrieved full texts, assessed study resources, and extracted data according to inclusion and exclusion criteria. The extracted data included first author, year of publication, sample size, source of miRNAs (plasma, serum, blood), number of analyzed miRNAs, type of data (raw expression or DE with fold-changes and P-values), type of DE analysis, and raw expression data/DE analysis results for all analyzed miRNAs. A total of 41 (0.2%) rows with missing and erroneous values in the input data were removed prior to meta-analysis. No missing values imputation was performed. To limit the bias from random one-off effect, only miRNAs identified by at least three independently reported studies were included in the analyses.

### Quality assessment

As recommended by the Cochrane Collaboration^23^, all studies selected for this analysis underwent quality assessment. The quality was assessed using the Newcastle-Ottawa Scale, which evaluates selection of study groups, comparability of groups and assessment of outcomes^24^. In addition, vote-counts of the general trend in directionality of miRNA changes in AD patients compared with healthy subjects were calculated by establishing the level of concordance in directionality of individual miRNA expression changes (up- or down-regulation) reported in different studies.

### Statistical analysis

All statistical analyses were performed in RStudio (v.2024.04.2 build 764, with R environment v.4.4.0). All P-values <0.05 or -log_10_ P-values >1.301 were considered significant. DE results for studies with raw expression data were calculated using deseq2 and limma packages or using the Wilcox test with the Benjamini-Hochberg correction. If statistical method was described in the original study, we used the same procedure in our analysis. Differences between groups of count values were analyzed using Pearson’s chi-square test. The conditions for the applicability of the statistical tests (such as normality of distribution, etc.) were verified prior to the analysis.

### Fold-change-based meta-analysis

In the meta-analysis, the effects of individual plasma miRNAs in AD patients compared with healthy subjects was examined using the Amanida package for R^25^. This method provided for each miRNA information about the directionality of miRNA expression change (up- or down-regulated), compound log_2_ fold-change and compound P-value. Compound log_2_ fold-change was calculated as the average of the individual log-transformed (base 2) fold-changes from the input studies weighted by study sizes. The compound P-value was calculated using a Fisher-based weighted P-value combination of the individual P-values from the input studies weighted by the study size.

In addition, pseudo-T-scores were calculated using the formula *[(avg (logFC) / sd (logFC) * sqrt (N_studies_)]*^26^. This score measures consistency in directionality in miRNA expression changes, with absolute values above 1.96 indicating the highest consistency in up- or down-regulated miRNAs across the studies. The compound P-values and the pseudo-T-scores were used in combination to identify the most significant and consistent miRNA changes in AD patients compared with healthy controls across the studies.

### Weighted miRNAs co-expression network meta-analysis

Clusters of similarly behaving miRNA profiles in AD patients were identified using a weighted miRNA co-expression network analysis (WmiRNACNA). This analysis was based on the general workflow derived from the weighted gene co-expression network analysis (WGCNA) but with the input represented by the log_2_ fold-change values from individual studies describing differences in miRNAs expression in AD patients compared with healthy subjects. A weighted adjacency matrix was computed by calculating the signed biweight midcorrelation *[cor_i,j_ = (1 + bicor) / 2]* between all miRNAs. Due to different number of studies for each miRNA, ’pairwise.complete.obs’ command was used. Optimal soft-threshold 18 was next calculated using the WGCNA::pickSoftThreshold.fromSimilarity function (with a resulting R^2^=0.868) and the correlation matrix raised to this power. The signed Topological Overlap Matrix (TOM) was obtained using the WGCNA::TOMsimilarity function and acceptable scale-free network properties of the matrix verified. This matrix was converted to a dissimilarity matrix using 1-TOM. Last, a hierarchical cluster analysis using Ward’s D^2^ method was performed. The obtained dendrogram was partitioned into individual clusters using the function WGCNA::cutreeDynamic (method=’hybrid’, deepSplit=2, pamStage=T, pamRespectsDendro=T, minClusterSize=10). Subsequently, similar clusters were merged using the WGCNA::mergeCloseModules function (with Module Eigengene dissimilarity threshold of 0.4).

In traditional WGCNA, clusters of interest are identified by correlating kME values (eigengenes of modules) with defined phenotypic information about compared samples, for example, AD patients versus healthy subjects. Since our data do not allow direct comparison between AD patients and healthy subjects in terms of similarly behaving miRNAs, AD-related clusters were identified using the following criteria: (1) cluster miRNAs’ log_2_ fold-change values were significantly different from zero based on one-sample Mann-Whitney-Wilcox test, (2) cluster miRNAs’ log_2_FC Q1-Q3 range was outside of zero and, (3) at least 50% of miRNAs in a cluster were significantly different in AD compared with healthy subjects according to the Amanida-based meta-analysis.

### MiRNA targets prediction

Prediction of the miRNA targets was performed by miRTargetLink 2.0 (https://ccb-compute.cs.uni-saarland.de/mirtargetlink2/), which works with validated (miRTarBase 8.0) and predicted (mirDIP, miRDB) miRNA targets from Homo sapiens^27^. The final set of target genes corresponding to significantly changed miRNAs in AD compared with healthy subjects was established only after removal of duplicate results. MiRNAs were classified as **“**AD-known**”** if they had at least one experimentally validated target (miRTarBase 8.0) recognized to be directly implicated in AD pathogenesis (e.g., APP, ADAM10, BACE1, PSEN1/2, MAPT). MiRNAs without validated interactions with AD-related genes were labeled **“**AD-unknown**.”** This division into „AD-known“ and „AD-unknown“ miRNAs was used only for data interpretation purposes and did not affect any statistical analyses.

### Functional enrichment analysis

Functional enrichment analysis of sets of genes targeted by selected miRNAs was performed using the ShinyGO V0.80^28^. Benjamini-Hochberg FDR correction was used to assess the significance level; 20 genes were selected as the minimum threshold for pathway size. MiRNA target functions were obtained using either REACTOME, Panther or KEGG database.

### Tissue specificity and involvement in AD

TissueAtlas version 2025 was used to analyze miRNA tissue specificity^29^. The presence of miRNAs in individual tissues was verified against a matrix of average miRNA expressions from TA. Only miRNAs with average tissue expression of ≥ 10 rpmm were included in the analyses. MiRNAs with zero expression were excluded prior to analysis.

### MiRNA coding motifs

To test what proportion of miRNAs originates from EVs, we calculated percentages of plasma miRNAs corresponding to miRNAs previously identified in EVs as annotated in Vesiclepedia, a manually curated database of molecules identified in different classes of EVs^30^. To find out proportions of miRNAs harbouring EXO motifs^3^, we first established which EXO/CELL motifs are present in each miRNA by cross-referencing the sequences of individual motifs against the miRNA sequences. Only complete EXO/CELL sequence in the miRNA was considered as the presence of a motif. We then calculated the sum of EXO and CELL motifs per miRNA. MiRNAs were considered carrying EXO motifs only if their miRNA sequence contained exclusively EXO motifs or the number of EXO motifs in the sequence was greater than the number of cellular retention CELL motifs.

## Results

### Sample quality and characteristics

A search using predefined key words identified 762 publications in PubMed/PMC and 73 GEO repository datasets (Extended Data Figure 1). Abstract review excluded 555 records, because studies reported animal research, lacked original research or presented meta-analyses. Full text review excluded further 258 records reporting miRNAs analyses from sources other than plasma, serum or blood, without full-text or showing incomplete results or incomplete data. Following review of individual miRNAs reported in the remaining 22 records, an additional 245 miRNAs were excluded due to questionable reproducibility as they were not detected by more than two independent studies. The study sample passed quality control based on the Newcastle-Ottawa scale (Extended Data Figure 2). Since 82% of miRNAs showed the same direction of expression change in at least 75% of studies, no miRNAs were excluded based on vote counts (Extended Data Figure 3, Extended Data Table 3). The final study sample consisted of 2650 miRNAs in a cohort of 4186 individuals (Table 1)^21,31–49^.

**Table 1.**
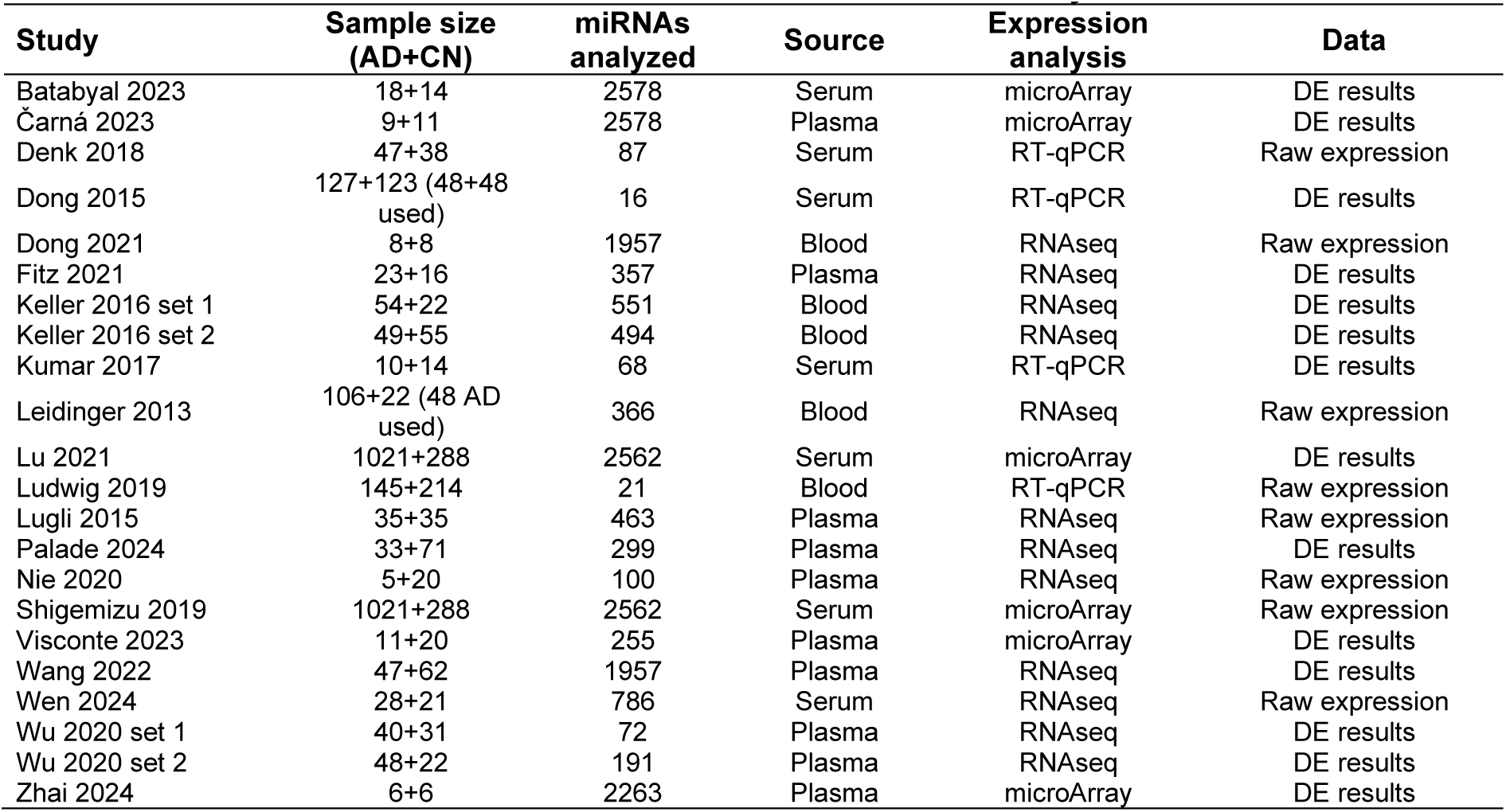
Main characteristics of studies included to meta-analysis.

| Study | Sample size<br>(AD+CN) | miRNAs<br>analyzed | Source | Expression<br>analysis | Data |
| --- | --- | --- | --- | --- | --- |
| Batabyal 2023 | 18+14 | 2578 | Serum | microArray | DE results |
| Čarná 2023 | 9+11 | 2578 | Plasma | microArray | DE results |
| Denk 2018 | 47+38 | 87 | Serum | RT-qPCR | Raw expression |
| Dong 2015 | 127+123 (48+48<br>used) | 16 | Serum | RT-qPCR | DE results |
| Dong 2021 | 8+8 | 1957 | Blood | RNAseq | Raw expression |
| Fitz 2021 | 23+16 | 357 | Plasma | RNAseq | DE results |
| Keller 2016 set 1 | 54+22 | 551 | Blood | RNAseq | DE results |
| Keller 2016 set 2 | 49+55 | 494 | Blood | RNAseq | DE results |
| Kumar 2017 | 10+14 | 68 | Serum | RT-qPCR | DE results |
| Leidinger 2013 | 106+22 (48 AD<br>used) | 366 | Blood | RNAseq | Raw expression |
| Lu 2021 | 1021+288 | 2562 | Serum | microArray | DE results |
| Ludwig 2019 | 145+214 | 21 | Blood | RT-qPCR | Raw expression |
| Lugli 2015 | 35+35 | 463 | Plasma | RNAseq | Raw expression |
| Palade 2024 | 33+71 | 299 | Plasma | RNAseq | DE results |
| Nie 2020 | 5+20 | 100 | Plasma | RNAseq | Raw expression |
| Shigemizu 2019 | 1021+288 | 2562 | Serum | microArray | Raw expression |
| Visconte 2023 | 11+20 | 255 | Plasma | microArray | DE results |
| Wang 2022 | 47+62 | 1957 | Plasma | RNAseq | DE results |
| Wen 2024 | 28+21 | 786 | Serum | RNAseq | Raw expression |
| Wu 2020 set 1 | 40+31 | 72 | Plasma | RNAseq | DE results |
| Wu 2020 set 2 | 48+22 | 191 | Plasma | RNAseq | DE results |
| Zhai 2024 | 6+6 | 2263 | Plasma | microArray | DE results |

### Meta-analysis of circulating miRNAs in AD

To obtain a comprehensive understanding of circulating miRNAs in AD, we measured their expression in AD patients and healthy subjects. A total of 910 out of 2650 miRNAs demonstrated significant expression changes in AD (compound P-scores, Figure 1A, Supplementary Table 1A). Only 194 of these miRNAs, however, showed the same direction of expression change (pseudo-T-scores, i.e. always up- or down-regulated) in at least 3 independent studies. 161 of these miRNAs were up- and 33 down-regulated in AD (Extended Data Figure 4, Extended Data Table 4). Targets of these “AD miRNAs” were projected to dysregulate cytokine and receptor tyrosine kinase (TRK) signaling, to repress cell death and estrogen-receptor (ER) pathways and to accentuate signal transduction and transcription (Extended Data Figure 5, Extended Data Table 5).

**Figure 1.**
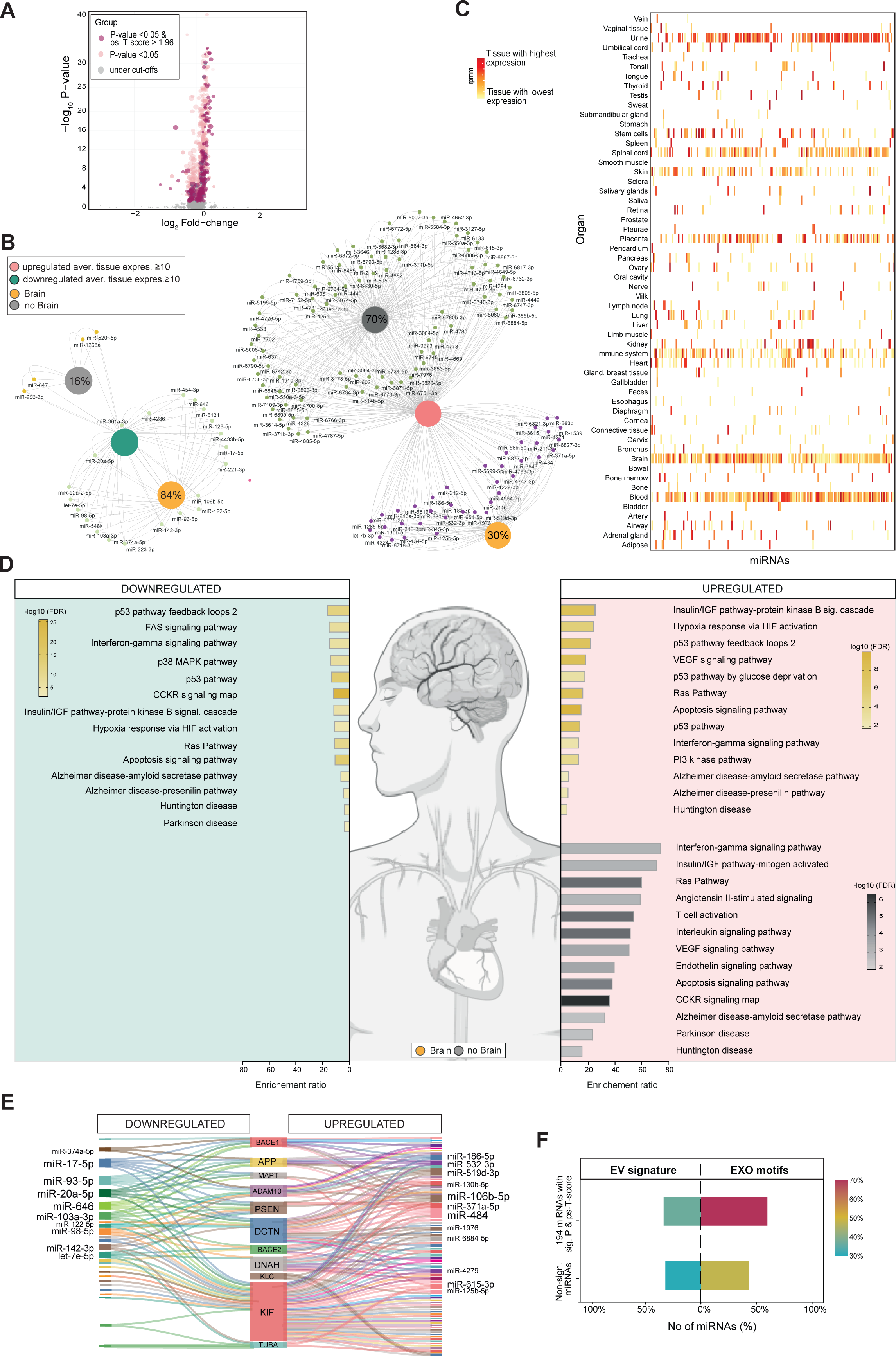
Amanida-based meta-analysis. (**a**) Volcano plot showing Amanida-derived meta-analysis results. Red circles correspond to miRNAs with compound P-values -log_10_ >1.301 and pseudo-T-scores >1.96, yellow and grey circles correspond to miRNAs with compound P-values -log_10_ > or < 1.301, respectively. (**b**) Venn networks showing the prevalence of significantly up- (pink) and down- (green) regulated miRNAs described (yellow) or not (grey) in the brain in AD patients compared with healthy subjects (average expression level ≥ 10 rpmm). (**c**) Presence of “AD miRNAs” in individual tissues based on the miRNATissueAtlas version 2025 (one miRNA was not present in the TissueAtlas database). Only 10 tissues with the highest average miRNA expressions (rpmm normalized counts) are displayed for each miRNA changed in AD. (**d**) The top 10 most significantly enriched and top neurodegeneration-related PANTHER knowledgebase forecast biological pathways based on the strong targets of significantly up-and down-regulated miRNAs found in brain or other tissues in AD patients compared with healthy subjects. (**e**) Sankey plot showing targets known to be involved in the pathogenesis of AD including targets involved in impaired axonal transport for up- (pink) and down- (green) regulated miRNAs in AD patients compared with healthy subjects. MiRNAs with more than 3 related AD targets are displayed in the left panel. (**f**) Combined barplot of Vesiclepedia-predicted proportions of miRNAs found in extracellular vesicles (EVs) (P=0.708) and percentage of secretion-promoting EXO motifs (P=0.00002) in AD patients compared with healthy subjects.

We next investigated possible tissues of origin of these “AD miRNAs”. 30 and 84% of up- and down-regulated “AD miRNAs”, respectively, were previously described in the brain but concomitantly found most also in other organs (Figure 1B and C, Supplementary Table 1B and 1C). Key targets of brain “AD miRNAs” were predicted to dysregulate p53, insulin/IGF, Ras and interferon-γ signaling, hypoxia-response via HIF activation, and apoptosis as well as AD and Huntington’s disease (HD) pathways, to accentuate CCKR, FAS and p38 MAPK signaling and Parkinson’s disease (PD) pathways, and to reduce VEGF and PI3 kinase signaling (Figure 1D, Supplementary Table 1D). Key targets of “AD miRNA” found likely to originate from outside the brain were predicted to equally dysregulate insulin/IGF, Ras and interferon-γ signaling, apoptosis and AD, HD and PD pathways and to reduce VEGF, CCKR, angiotensin II, interleukin, endothelin signaling and T cell activation. Apart from Apolipoprotein E, many molecules linked to the pathogenesis of AD including APP and tau, as well as components of axonal transport, were all found among “AD miRNA” targets (Figure 1E, Supplementary Table 1E). Independently from their tissues of origin, miRNAs were projected to be enriched in EXO motifs in AD (Figure 1F, Supplementary Table 1F).

### WmiRNACNA in AD

To identify networks of miRNAs that are changed in AD, we developed WmiRNACNA by modifying WGCNA (in Methods). Based on log_2_ fold-changes, hierarchical clustering of 2650 miRNAs gave rise to 12 miRNA membership modules (Figure 2A, Supplementary Table 2A). Modules E, F and K showed average log_2_ fold-change values furthest from zero (Figure 2B, Supplementary Table 2B), and contained the highest percentages of “AD miRNAs” previously found by meta-analysis (Figure 2C, Supplementary Table 2C). All miRNAs in these modules were upregulated. Key targets of “AD module miRNAs” were predicted to impact the same activities as targets of “AD miRNAs” identified by meta-analysis including cytokine (E and F), RTK and estrogen signaling (F) in addition to cell cycle regulation (E and K, Extended Data Figure 6, Extended Data Table 6).

**Figure 2.**
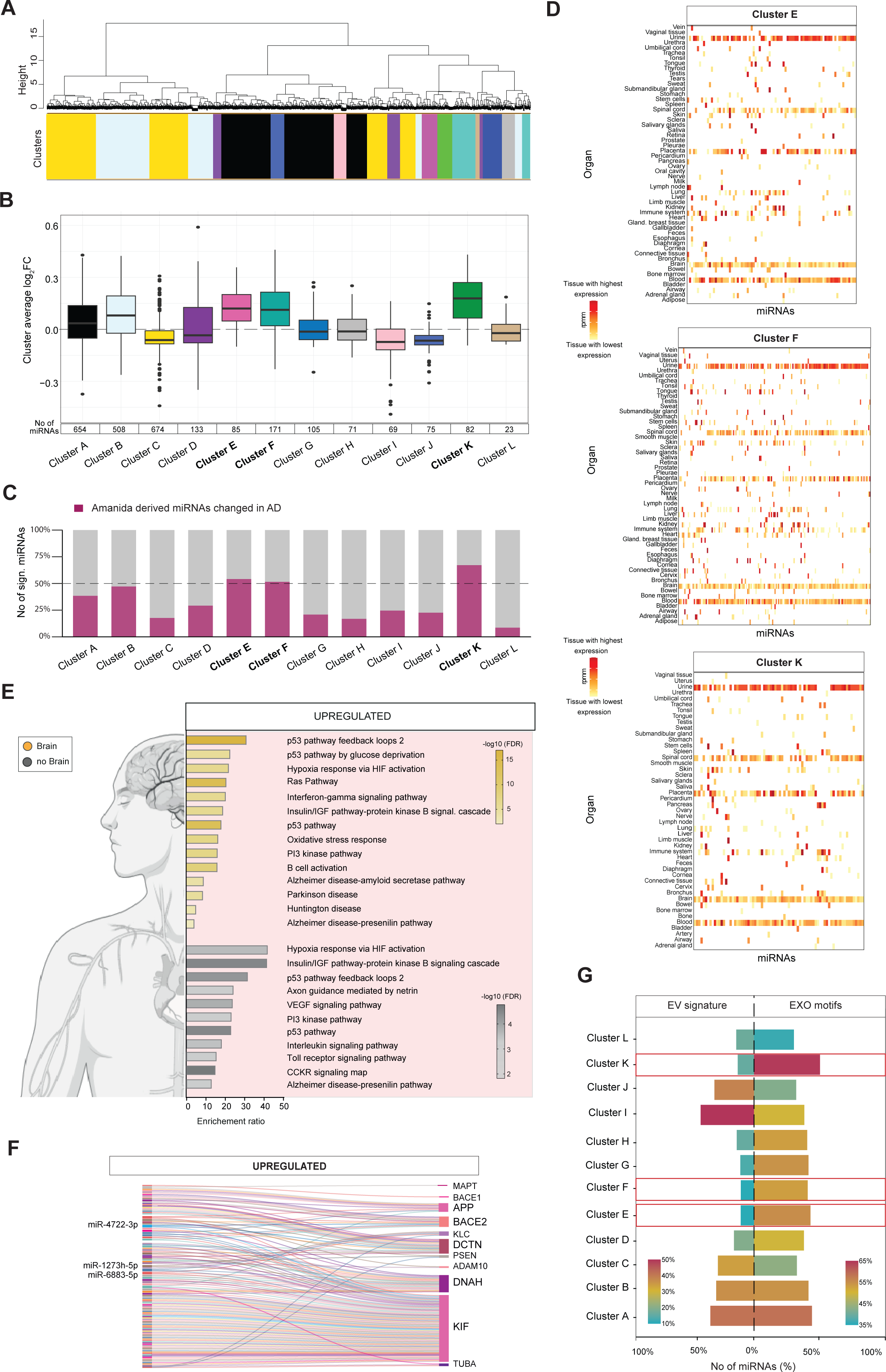
Weighted miRNAs co-expression network analysis. (**a**) Dendrogram depicting results of hierarchical clustering, bottom color strip indicates distribution of the final 12 modules. (**b**) Boxplots portraying distribution of average log_2_FC values (calculated as mean of log2FC values from individual studies included in meta-analysis) of miRNAs in individual modules. Bottom table shows number of miRNAs in each module. Modules E, F, and K (depicted in bold) were characterized by the most significantly changed log2FC values (Q1–Q3 ≠ 0) and contained the highest proportion of miRNAs significantly altered in AD as revealed by the meta-analysis. (**c**) Barplot showing proportion of miRNAs changed in AD patients based on meta-analysis in individual modules. E, F and K modules with most significantly increased proportions of changed miRNAs in AD patients are emphasized in bold. Modules C, G, H and L exhibit least miRNA changes. (**d**) Heatmaps showing presence of miRNAs from modules E, F and K in individual tissues (85[100%], 162 [95%] and 75 [93%]) of miRNAs were represented in the database). Only 10 tissues with the highest average miRNA expressions (rpmm normalized counts) are displayed for each miRNA changed in AD. (**e**) The top 10 most significantly enriched and neurodegeneration-related PANTHER knowledgebase forecast biological pathways based on targets of miRNAs found inside or outside the brain in modules E, F, and K. (**f**) Sankey diagram showing targets known to be involved in the pathogenesis of AD. MiRNAs with more than 3 related AD targets are displayed in the left panel. (**g**) Combined barplot of Vesiclepedia-based predicted proportions of miRNAs found in extracellular vesicles (EVs) and harboring secretion-promoting EXO motifs in WmiRNACNA clusters. Red frames indicate AD-related modules.

We next asked whether miRNAs in “AD modules” are found predominantly in the brain. Only 21, 21 and 25% of E, F, and K module miRNAs were previously described in the brain (Figure 2D, Supplementary Table 2D, Extended Data Table 2). The most frequent targets of these “AD module miRNAs” were projected to suppress p53, insulin/IGF, Ras, interferon-γ and PI3 kinase signaling, hypoxia-response via HIF activation, oxidative stress, B cell activation as well as AD pathways (Figure 2E, Supplementary Table 2E). “AD module miRNAs” found outside the brain were projected to reduce interleukin signaling and Rho GTPase cytoskeletal regulation. Apart from Apolipoprotein E, all molecules linked to the pathogenesis of AD including APP and tau as well as components of the axonal transport, were found among “AD miRNA module” targets (Figure 2F, Supplementary Table 2F). Projected frequency of miRNAs in EVs as well as enriched in EXO motifs varied significantly between modules (Figure 2G, Supplementary Table 2G).

### Key circulating miRNAs in AD

A total of 37 miRNAs were found significantly changed in AD based on both meta-analysis and the WmiRNACNA (Extended Data Figure 7, Table 2, Extended Data Table 7). Only 7 of these miRNAs were previously linked to AD with targets including APP, ADAM10, BACE, and presenilin (Figure 3A, Supplementary Table 3A). Targets of these “AD miRNAs” were predicted to suppress axon guidance, p53 pathway, Wnt and cadherin signaling, angiogenesis and Alzheimer’s disease-presenilin pathways (Figure 3B, Supplementary Table 3B). The other 30 miRNAs remain poorly described in the context of AD. Their targets are projected to center frequently around the CCKR signaling map and to most commonly dysregulate insulin/IGF, p38 MAPK, Toll-receptor, PDGF and the interleukin signaling pathways. The “AD miRNAs” previously linked to AD showed the highest levels of expression in liver and immune system related tissues, while poorly known miRNAs in AD were found primarily in blood and the vestibulo-cochlear complex with several orders of magnitude lower expression levels (difference 10^4^ rpmm, Figure 3C, Supplementary Table 3C). We last attempted to understand functional repercussions of changes in these key circulating “AD miRNAs”. Based on Kyoto Encyclopedia of Genes and Genomes, all “AD miRNAs” were predicted to play a dominant role in axon guidance and cancer pathways (Figure 3D, Supplementary Table 3D). In addition, “AD miRNAs” previously linked to AD were predicted to contribute to oxytocin signaling, adherent junctions, TGFβ signaling, longevity, hedgehog signaling, neurotransmitter signaling and AD and neurodegeneration, while poorly described “AD miRNAs” in AD were predicted to play roles in ErbB, hippo and p53 signaling.

**Figure 3.**
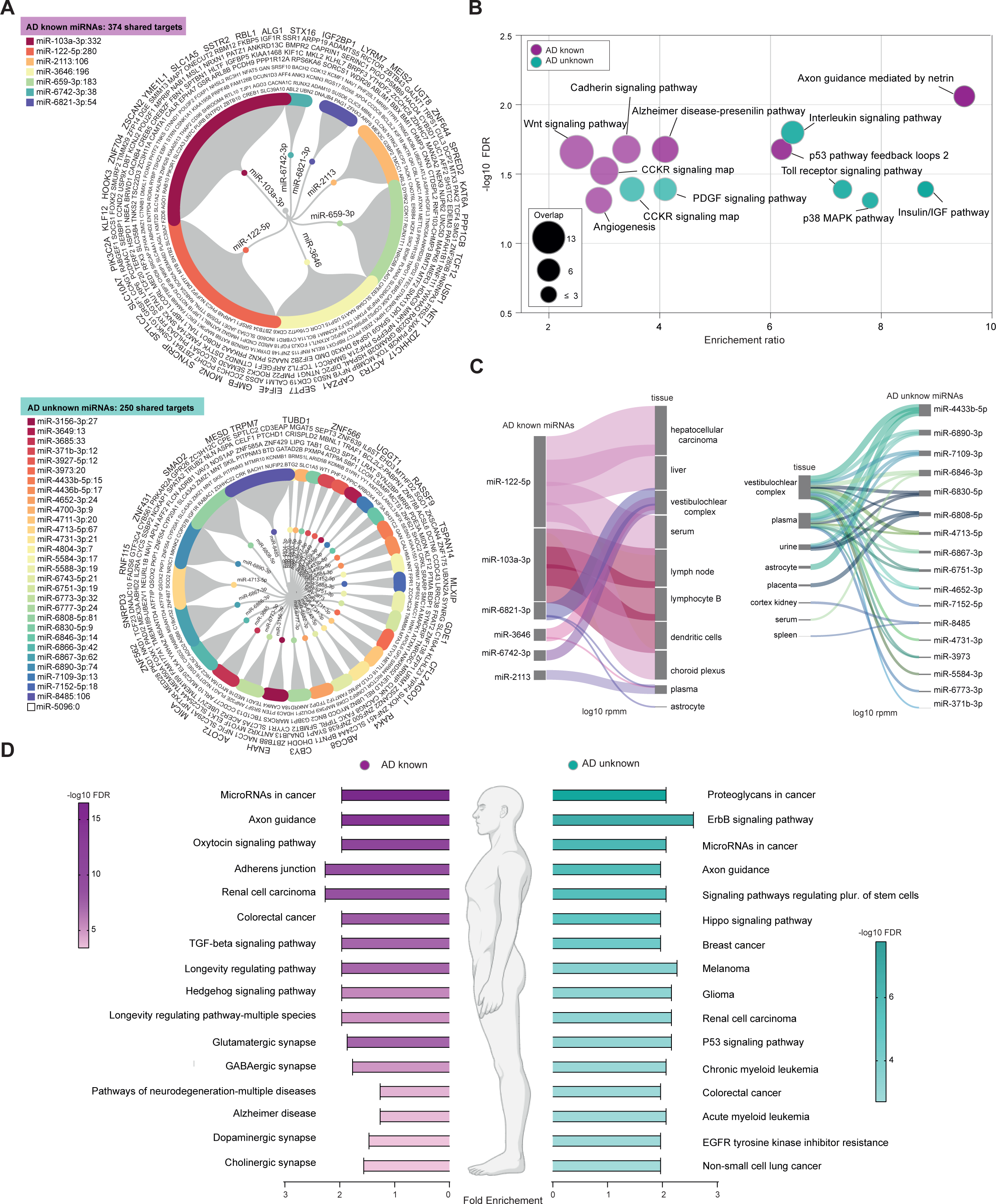
Key AD-related miRNAs. (**a**) Number of miRNAs identified as known or unknown to play roles in AD pathogenesis based on their targets. Circular dendrograms show proportion of targets involved in biological processes of miRNAs known or unknown to play roles in AD pathogenesis. Minimum shared target was set to 3. (**b**) Most significant PANTHER knowledgebase forecast biological pathways identified for the minimum of 3 shared targets of AD known and unknown miRNAs. (**c**) Sankey plot showing tissue specificity for AD known and unknown miRNAs. The top 3 individual tissues per miRNA based on miRNATissueAtlas version 2025 database are shown (average expression level ≥10 rpmm). Line thickness represents the miRNA expression level in individual tissue (log10(rpmm)=1to6). (**d**) Top 10 complex interactions and disease related pathways (KEGG) for AD known and AD unknown miRNAs.

**Table 2.**
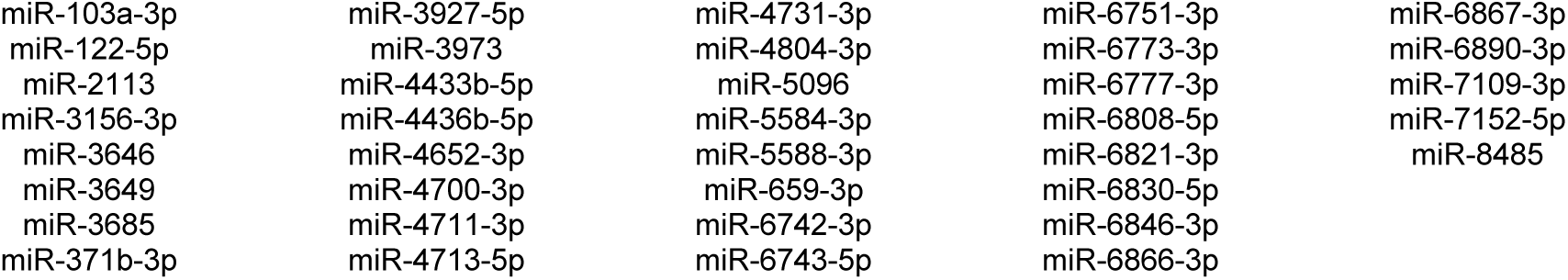
The most relevant AD-related miRNAs derived from both meta-analyses.

|  |  |  |  |  |
| --- | --- | --- | --- | --- |
| miR-103a-3p | miR-3927-5p | miR-4731-3p | miR-6751-3p | miR-6867-3p |
| miR-122-5p | miR-3973 | miR-4804-3p | miR-6773-3p | miR-6890-3p |
| miR-2113 | miR-4433b-5p | miR-5096 | miR-6777-3p | miR-7109-3p |
| miR-3156-3p | miR-4436b-5p | miR-5584-3p | miR-6808-5p | miR-7152-5p |
| miR-3646 | miR-4652-3p | miR-5588-3p | miR-6821-3p | miR-8485 |
| miR-3649 | miR-4700-3p | miR-659-3p | miR-6830-5p |  |
| miR-3685 | miR-4711-3p | miR-6742-3p | miR-6846-3p |  |
| miR-371b-3p | miR-4713-5p | miR-6743-5p | miR-6866-3p |  |

## Discussion

A major strength of this study is the rigor with which data have been collected and analyzed. Given that many studies reporting miRNA abnormalities in AD provide only data for significantly DE miRNAs, there is a risk that the observed abnormalities in individual miRNAs are biased by overestimating their significance given the lack of data on instances where the miRNAs in question did not show significant differences. To circumvent this potential bias, this meta-analysis has investigated the pooled effect of miRNA abnormalities in AD based only on studies that provided a complete set of results either from DE analyses which include also non-significantly DE miRNAs, or from a complete list of raw miRNA expression data. This study used different meta-analytic approaches, analyzing first the individual roles of the most significant miRNAs in AD and then identifying modules of miRNAs exhibiting similar behavior in AD. This multivariate approach highlights key “AD miRNAs” and contributes to understanding AD better. One potential limitation of the study is the inconsistency of naming miRNAs in the original studies. Given the evolution of miRNA nomenclature over time, for example the use of * and 3p/5p refinements, the names of miRNAs in different studies may differ, although sequence-wise they refer to the same miRNA. This issue, however, cannot be resolved within the secondary analysis without full access to sequence data and should therefore be kept in mind when interpreting our results. In the future, it would be useful to implement tools for retrospective application of currently valid miRNA nomenclature standards to earlier studies.

Using two meta-analyses, we identified a number of “AD miRNAs”. Surprisingly, many of these miRNAs have not yet been characterized as AD-related, so their identification broadens our horizons and opportunities in exploring the mechanisms of AD and the role of these miRNAs. Several molecules previously linked to the pathogenesis of AD have been identified among targets of these “AD miRNAs”. Observation that miRNAs target the entire Aβ pathway implicates circulating miRNAs directly in the pathogenesis of AD. Molecules involved in impaired axonal transport in AD including molecular motors, besides microtubule interacting proteins such as tau, have also been found among targets of “AD miRNAs,” further supporting a role of miRNAs in AD. Altogether, these results provide a comprehensive understanding of the contribution of circulating miRNAs in the pathogenesis of AD and, in addition, corroborate previously noted associations between miRNAs and AD molecules^50^. Studies of mechanisms underlying miRNA changes offer some evidence of dysregulated Drosha^51^ and Dicer^52^. Alternatively, given that most “AD miRNAs” are enriched in EXO motifs and therefore preferentially secreted, it might well be that miRNA defects occur at the level of the endosomal/lysosomal pathway^53,54^. Significant further work is needed to establish the mechanisms and pinpoint origins underlying systemic circulating miRNA changes in AD. Targets of “AD miRNAs” reveal involvement of most diverse pathways in the pathogenesis of AD. Some of the targets are involved in the cell cycle and death^55^ as well as in cytokine^56^, estrogen^57^ and insulin^58–60^ signaling, which have all been previously described in AD. Other targets, for example CCKR^61–63^, integrin^64^, interferon-γ^65^ and p53^66,67^ signaling as well as angiogenesis^68^ have hardly been accounted for in the context of AD. These pathways represent knowledge gaps in our understanding of AD. For example, miRNAs projected to dysregulate p53 in AD might provide valuable clues in understanding the interactions between cellular senescence, AD and cancer. Surprisingly, only a fraction of “AD miRNAs” has been described in the brain and at the same time, most frequently as relevant in several other tissues. This suggests that “AD miRNAs” belonging to different tissues of origin are functionally diverse in AD. This is unexpected, considering AD is thought to be a neurodegenerative disorder with etiology exclusive to the brain. There are at least two plausible explanations of this finding. First, AD pathology triggers initial miRNA changes in the brain and only later in other tissues. And second, “AD miRNAs” from other tissues are generated independently from the brain and contribute to AD pathology. Both scenarios provide support to the view that AD is a systemic disorder. In conclusion, the findings presented in this study reveal novel circulating miRNAs altered in AD that functionally not only recapitulate most well-established pathways but also uncover several unaccounted-for pathways in AD and are thus most informative about its pathogenesis. These findings offer an unprecedented opportunity to develop a next generation of biomarkers, and open novel avenues for design of AD therapies.

## Statements

### Funding

The study was funded by the European Union: Next Generation EU – Project National Institute for Neurological Research (LX22NPO5107 (MEYS)).

### Competing Interests Statement

No disclosures.

### Open Science Statement

In this study, we adhere to the principles of Open Science as much as possible: we explicitly labelled the study as meta-analysis, we report the source studies used in meta-analysis, the data are available within the paper and its supplementary materials, the R script of the analysis is available on request from the first author, and all materials used are cited.

### Data availability statement

The data used in this study are available within the article and its supplementary materials.

### Authors contribution

The authors contributed to this article as follows: Conceptualization: GBS, Methodology: JSN, MC Investigation: JSN, Data curation: JSN, MC, Formal analysis: JSN, MC, Funding acquisition: GBS, Visualization: JSN, MC, Writing - original draft: JSN, MC, GBS, Writing - Review & Editing: EBD, ZM.

### Authors’ Note

No humans or animal models were used in our research.

## Extended Data Figure legends

**Extended Data Figure 1 - Systematic search of the database to identify circulating miRNAs for the meta-analyses.**

Flowchart of the selection process for the studies of plasma, serum and blood miRNAs changed in AD patients compared with healthy subjects to be included in the analysis.

**Extended Data Figure 2 - Newcastle-Ottawa Scale-based analysis.**

Newcastle-Ottawa Scale-based analysis of the quality of included studies.

**Extended Data Figure 3 - Barplot showing the rate of vote agreement.**

Stability of the deregulation trend of 194 AD-related miRNAs across studies included in the meta-analysis.

**Extended Data Figure 4 - Qualitative compound trend plot for 194 miRNAs with significant compound P-value and significant pseudo-T-score.**

Colored barplot shows number of studies in which miRNAs were up- and down-regulated.

**Extended Data Figure 5 – Meta-analysis based biological pathways changed in AD.** The top 5 most significantly enriched Reactome database forecast biological pathways based on the targets of significantly up- (pink circles) and down- (green circles) regulated miRNAs in AD patients compared with healthy subjects following meta-analysis.

**Extended Data Figure 6 – WmiRNACNA based biological pathway changes in AD.**

The top 3 most significantly enriched Reactome database forecast biological pathways based on the targets of significantly changes miRNAs in modules E, F and K.

**Extended Data Figure 7 – Key AD-related miRNAs.**

Venn diagram showing the intersection of the most significant AD-related miRNAs identified by Amanida-derived meta-analysis and the 3 most AD-related clusters containing miRNAs with similar behavior identified by weighted miRNAs co-expression meta-analysis.

## Notes

### Competing Interest Statement

The authors have declared no competing interest.

