## Supplementary figures and images for "Rethinking Alzheimer’s: Novel miRNAs Illuminate a Disease Beyond the Brain"

### Extended Data Figure 1

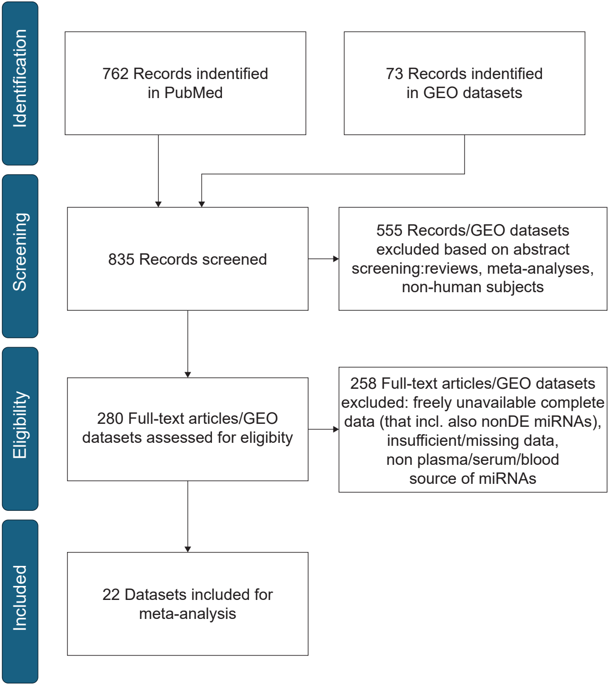

### Extended Data Figure 3

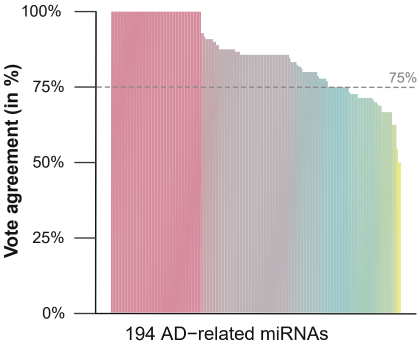

### Extended Data Figure 4

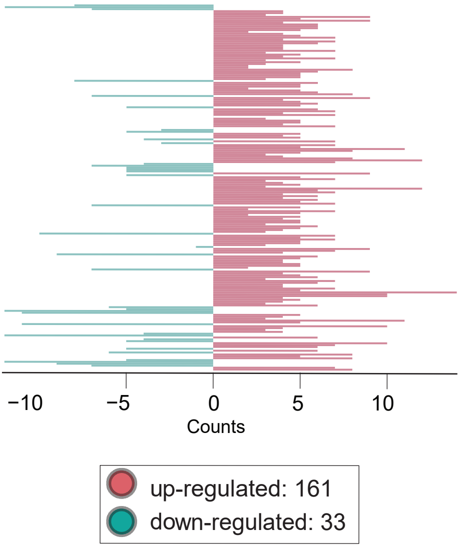

### Extended Data Figure 5

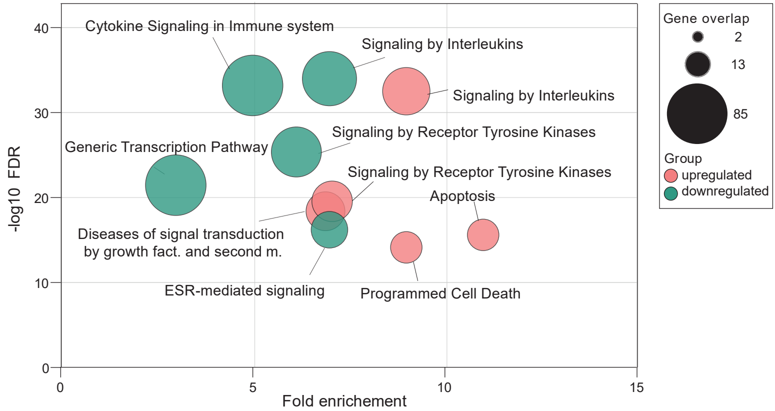

### Extended Data Figure 6

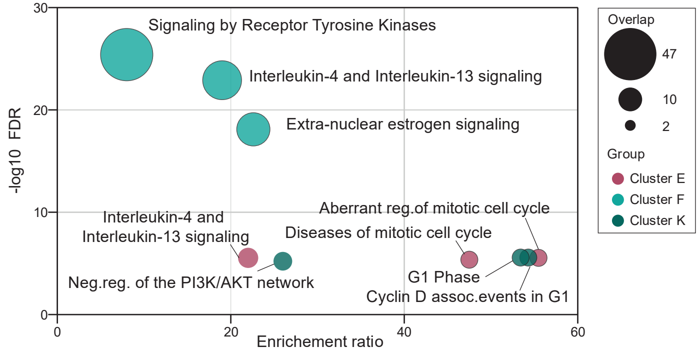

### Extended Data Figure 7

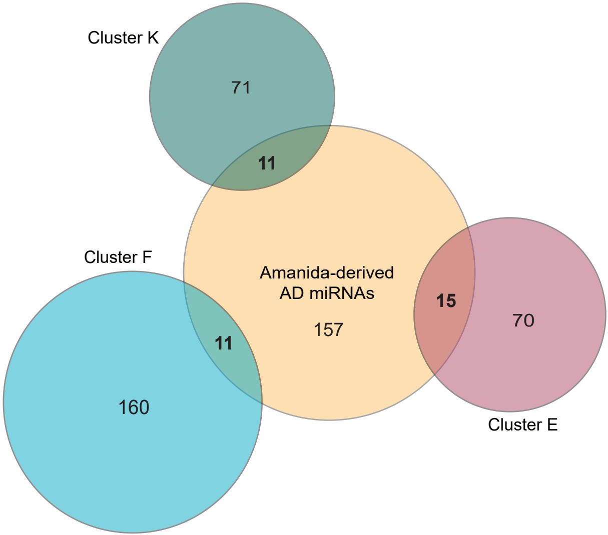
