## Extended Data Figure 2 for "Rethinking Alzheimer’s: Novel miRNAs Illuminate a Disease Beyond the Brain"

| | Selection of cases | | | Comparability | Measurement | |
|---|---|---|---|---|---|---|
| | AD cases definition | Representativness | Definition of healthy controls | | Measurement & analysis procedure | Same measurement for AD/CN |
| Batabyal 2023 | 1 | 1 | 1 | 1 | 1 | 1 |
| Čarná 2023 | 1 | 1 | 1 | 1 | 1 | 1 |
| Denk 2018 | 1 | 1 | 1 | 1 | 1 | 1 |
| Dong 2015 | 1 | 1 | 1 | 1 | 1 | 1 |
| Dong 2021 | 1 | 1 | 1 | 1 | 1 | 1 |
| Fitz 2021 | 1 | 1 | 1 | 1 | 1 | 1 |
| Keller 2016 | 1 | 1 | 1 | 1 | 1 | 1 |
| Kumar 2017 | 1 | 1 | 1 | 1 | 1 | 1 |
| Leidinger 2013 | 1 | 1 | 1 | 1 | 1 | 1 |
| Lu 2021 | 1 | 0 | 1 | 1 | 1 | 1 |
| Ludwig 2019 | 1 | 1 | 1 | 1 | 1 | 1 |
| Lugli 2015 | 1 | 1 | 1 | 1 | 1 | 1 |
| Nie 2020 | 1 | 1 | 0 | 0 | 1 | 1 |
| Palade 2024 | 1 | 1 | 1 | 1 | 1 | 1 |
| Shigemizu 2019 | 1 | 0 | 1 | 1 | 1 | 1 |
| Visconte 2023 | 1 | 1 | 1 | 1 | 1 | 1 |
| Wang 2022 | 1 | 1 | 1 | 1 | 1 | 1 |
| Wen 2024 | 1 | 1 | 1 | 1 | 1 | 1 |
| Wu 2020 | 1 | 1 | 1 | 1 | 1 | 1 |
| Zhai 2024 | 1 | 1 | 1 | 1 | 1 | 1 |
